# Adult sex ratio bias in snowy plovers is driven by sex-specific early survival: implications for mating systems and population growth

**DOI:** 10.1101/117580

**Authors:** Luke J. Eberhart-Phillips, Clemens Küpper, Tom E. X. Miller, Medardo Cruz-López, Kathryn H. Maher, Natalie dos Remedios, Martin A. Stoffel, Joseph I. Hoffman, Oliver Krüger, Tamás Székely

## Abstract

Adult sex ratio (ASR) is a central concept in population biology and a key factor in sexual selection, yet why do most demographic models ignore sex-biases? Vital rates often vary between the sexes and across life history, but their relative contributions to ASR variation remain poorly understood—an essential step to evaluate sex ratio theories in the wild and inform conservation. Here we combine structured two-sex population models with individual-based mark-recapture data from an intensively monitored polygamous population of snowy plovers. We show that a strongly male-biased ASR is primarily driven by sex-specific survival of juveniles, rather than adults or dependent offspring. This provides empirical support for theories of unbiased sex allocation when sex-differences in survival arise after the period of parental investment. Importantly, a conventional model ignoring sex-biases significantly overestimated population viability. We suggest that sex-specific population models are essential to understand the population dynamics of sexual organisms: reproduction and population growth is most sensitive to perturbations in survival of the limiting sex. Overall, our study suggests that sex-biased early survival may contribute towards mating system evolution and population persistence, with implications for both sexual selection theory and biodiversity conservation.

**SIGNIFICANCE STATEMENT:** Sex biases are widespread in nature and represent a fundamental component of sexual selection and population biology—but at which point in life history do these biases emerge? Here we report a detailed individual-based demographic analysis of an intensively studied wild bird population to evaluate the origins of sex biases and their consequences on mating strategies and population dynamics. We document a strongly male-biased adult sex ratio, which is consistent with behavioral observations of female-biased polygamy. Notably, sex-biased juvenile, rather than adult survival, contributed most to the adult sex ratio. Sex-biases also strongly influenced population viability, which was significantly overestimated when sex ratio and mating system were ignored. Our study therefore has implications for both sexual selection theory and biodiversity conservation.

---

Sex ratio variation in wild populations has important consequences for population dynamics and hence biodiversity conservation (1). As reproduction in sexual organisms involves both males and females, a shortage of either sex could compromise population viability (2). A reduction in the number of breeding females directly reduces birth rates and hence population productivity (3), whereas an overabundance of males may increase violence and aggression such that both male and female survival is reduced (4). Although a small number of males can potentially fertilize many mates, females in male-biased populations may need to compete for breeding opportunities with high quality males which can induce additional mortality (5). If males are in short supply, fathers also tend to reduce their parental investment which could negatively impact offspring survival (6, 7). Additionally, a biased sex ratio in either direction will decrease effective population size, which has adverse consequences for genetic diversity (8). Therefore, depending on mating system, populations with biased sex ratios may be more vulnerable to extinction than unbiased populations (9).

Recent studies also suggest that the adult sex ratio (ASR, the proportion of the adult population that is male) impacts breeding strategies as the limiting sex has the advantage in mating and parental decisions (10-12). For example, male-biased avian populations tend to have polyandrous mating systems and male-biased parental provisioning (13). Although the theory linking ASR to breeding system is relatively new, there are already supporting studies: parental cooperation is associated with an unbiased ASR in birds (14) whereas ASR is a strong predictor of sex-specific sexual activity and divorce rates in humans (15, 16).

Despite the importance of ASR in population biology, biodiversity conservation and breeding system evolution, the origin(s) of ASR biases remain unclear. Biases in the ASR can emerge via a number of mutually non-exclusive demographic pathways (11, 17, 18). For instance, sex-biases may occur at conception or at birth (19), or the survival of male and female juveniles may differ to the extent that fewer of one sex reaches adulthood (20).

Furthermore, sex-differences in adult survival or maturation rates could create a shortage of the sex that has higher mortality (4) or slower maturation (18), and if emigration is not compensated by immigration, sex-differences in dispersal behavior could create local biases in ASR (21).

A number of studies of wild vertebrate populations have evaluated the independent contributions of the above pathways to ASR bias (22-24). However, to fully understand ASR bias requires these components to be modeled simultaneously to quantify their relative contributions. In practice, large empirical datasets from natural populations incorporating stage- and sex-specific vital rates are uncommon (25-27). Furthermore, males and females often have different behaviors or ecological niches (28), which can make one or the other easier to detect (9, 11). Fortunately, these sources of sampling bias can be accounted for using mark-recapture methods (29).

Here, we investigate the demographic origins of ASR bias in a polygamous bird using seven years of individual-based sex- and stage-specific life history data. Polygamous species have a special significance in sex ratio studies as they are predicted to be at higher risk of extinction (1, 30). We studied a small ground-nesting shorebird, the snowy plover (*Charadrius nivosus*, ref. 31), which is endangered in parts of its Nearctic range and has a sequentially polygamous mating system (32, 33). Using a two-sex matrix model, we show that the ASR of this species is substantially more male biased than previously reported (34). Sex-differences in chick and juvenile survival contribute most to ASR bias, suggesting that ASR variation is particularly susceptible to factors that influence early life history stages. Furthermore, we show that population growth is most sensitive to adult female survival under a male-biased ASR, signifying that sex-specific early survival can affect population viability via ASR variation. Importantly, our study suggests that sex-biased survival in early life has ramifications for mating system variation and knock-on effects for population growth.

## RESULTS

We conducted this study at Bahía de Ceuta, a subtropical lagoon on the coastal plain in northwestern Mexico (23°54’N, 106°57’W). Between 2006 and 2012, we uniquely marked and monitored 1259 individuals (436 females and 390 males initially marked as chicks and 221 females and 212 males initially marked as adults). Although our marking methods were limited to breeding adults and chicks, we detected no sex difference in the proportion of this marked population that was non-breeding (paired t-test: *t* = 0.429, *df* = 4, *P* = 0.69, Fig. S1). Therefore, this marked subset of the population represents a broadly representative sample from which to draw inferences about the dynamics of the population at large and to elucidate the contributions of sex- and stage-specific survival towards ASR bias.

### Mating system

To understand the ASR in the context of mating system, we quantified sex-specific mating strategies of snowy plovers at our study site. Although both sexes can be polygamous, female snowy plovers typically desert broods to seek serial mates, leaving males to provide parental care alone (33). Thus, we expected that females would acquire on average a greater number of mates per year than males. Based on behavioral observations of 456 families with known identities of both parents, this is precisely what we found (Fig. 1; Mann-Whitney-Wilcoxon test: *W* = 3264, *P* ≤ 0.001). As such, the mating system index in the mating function (Eq. 4 in *Methods*) was polyandrous (*h* = 0.82).

**Figure 1.**
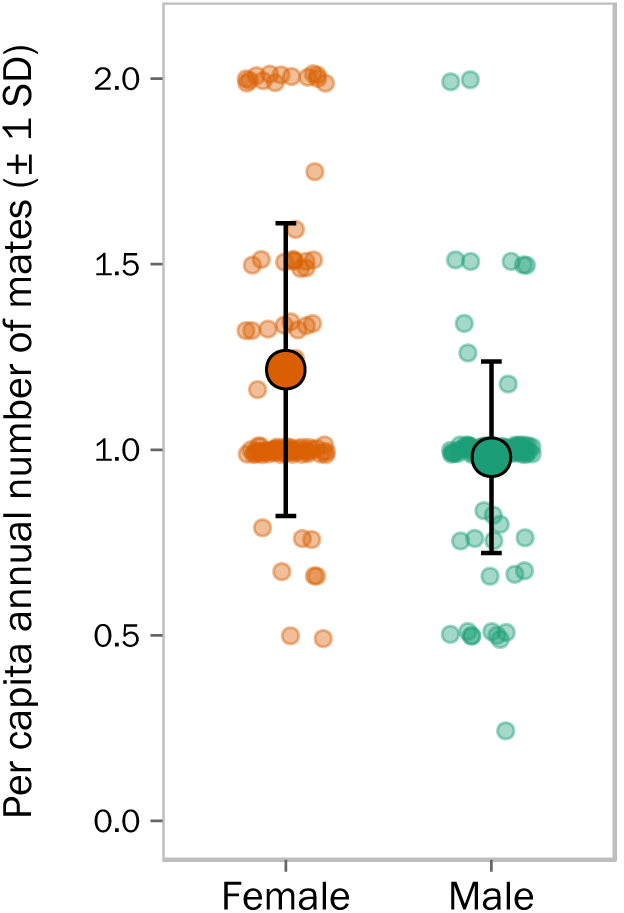
Sex bias in mating system illustrated as the per capita annual number of mates acquired by male and female snowy plovers (Mann-Whitney-Wilcoxon test: *W* = 3264, *P* < 0.001, *N* = 456 families).

### Sex-biased survival and ASR bias

We estimated stage- and sex-specific survival rates using mark-recapture analysis to control for imperfect detection in the field. Mark-recapture modeling revealed sex differences in encounter probability for juveniles and adults that would confound simple estimates of survival and ASR based solely on return rates or uncorrected counts of males and females (Table 1). Apparent survival was strongly male biased across all life-history stages, with male survival being 11.5% higher than female survival at the chick stage, 51% higher for juveniles, and 0.5% higher at the adult stage (Fig. 2a). Hatching sex ratio was slightly female biased but did not significantly deviate from parity (average *ρ* = 0.486 [95% CI = 0.435−0.536], *P* = 0.588, *N =* 340 hatchlings from 116 full broods). Overall, our model indicated a strongly male-biased ASR (mean = 0.632 [95% CI = 0.460−0.785]; Fig. 2b).

**Table 1.** Summary statistics of sex- and stage-specific estimates of the snowy plover population. Notation includes: *N* = number of individual encounter histories used for mark-recapture modelling, ϕ = apparent survival, and *p* = encounter probability. Estimates are shown as the median and 95% confidences interval of each bootstrapped distribution.

| Sex | Stage | $N$ | $\phi$ | $p$ |
| --- | --- | --- | --- | --- |
| Female | Chick | 416 | 0.48<br>(0.41–0.55) | 0.56<br>(0.53–0.58) |
|  | Juvenile | 234 | 0.15<br>(0.11–0.19) | 0.48<br>(0.34–0.62) |
|  | Adult | 221 | 0.68<br>(0.62–0.74) | 0.52<br>(0.40–0.65) |
| Male | Chick | 372 | 0.53<br>(0.47–0.60) | 0.55<br>(0.53–0.58) |
|  | Juvenile | 243 | 0.22<br>(0.18–0.27) | 0.66<br>(0.48–0.74) |
|  | Adult | 212 | 0.69<br>(0.63–0.74) | 0.66<br>(0.54–0.77) |

**Figure 2.**
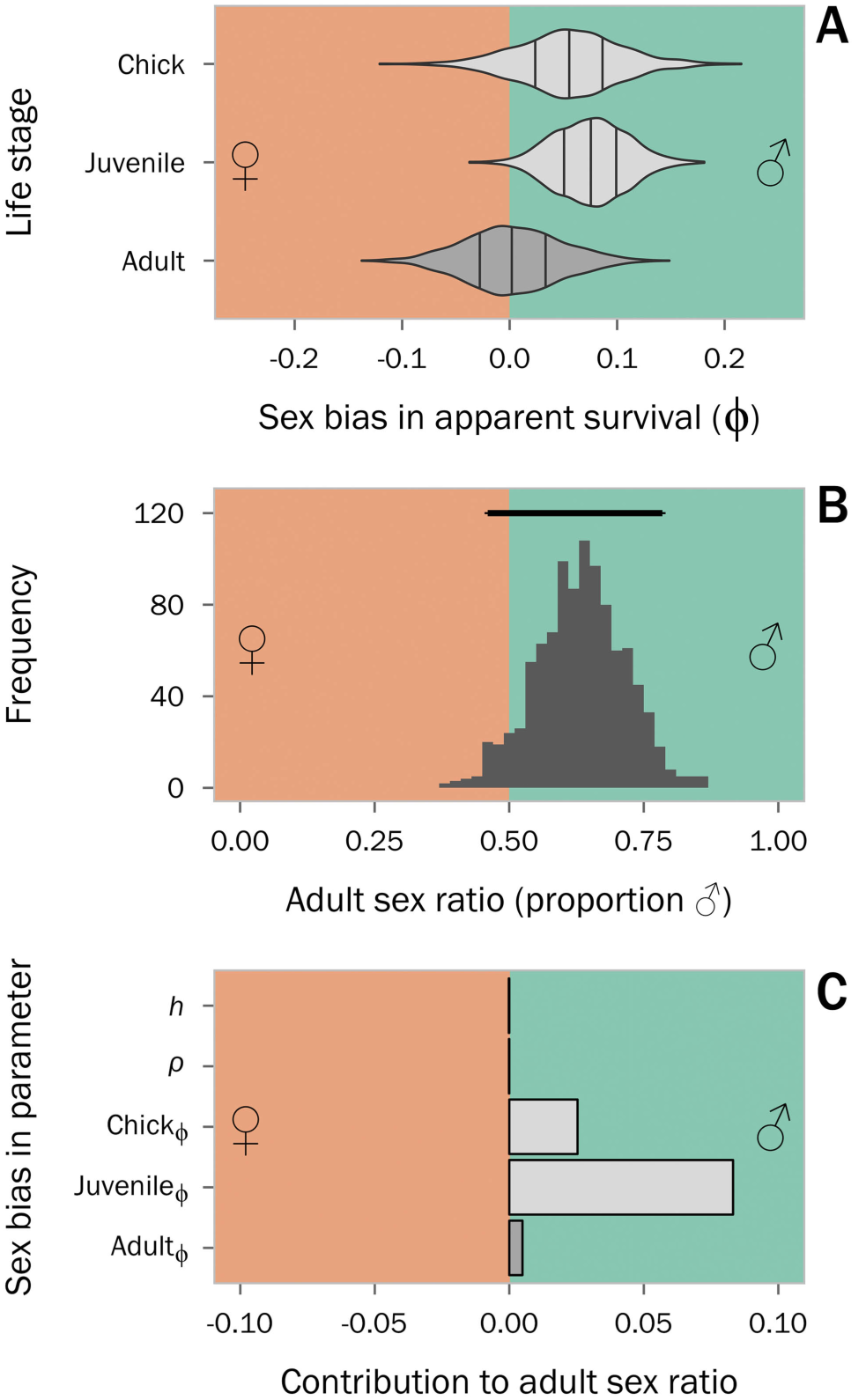
(a) Bootstrap distributions of stage-specific sex biases in apparent survival estimates, with internal lines indicating median and interquartile ranges, and shades of grey corresponding to first-year (light grey) or adult (dark grey) parameters shown in Fig. 6; (b) Bootstrap distribution of the derived ASR, with the horizontal bar above the distribution indicating the 95% confidence interval of the ASR estimate based on 1000 iterations (mean ASR = 0.632 [95% CI: 0.460 − 0.785]); (c) relative contributions of model components to ASR in our life table response experiment (LTRE) which compared the empirically-derived sex-specific model to a theoretical model in which survivorships did not differ between the sexes. Our measure of ASR was expressed as the proportion of the adult population that is male, thus changes in female-biased parameters have a negative effect on ASR and thus their LTRE statistics are negative.

### Contributions to male-biased ASR

To elucidate the stage-specific contributions of sex differences in survival to ASR bias, we conducted a life table response experiment (LTRE), which revealed that all vital rates contributed in the same direction (i.e. male-biased) but differed in magnitude. A sex difference in juvenile survival made the largest overall contribution to ASR bias (Fig. 2c). Specifically, the contribution of sex-biased juvenile survival towards ASR was 3.3 times higher than sex-biased chick survival and 17.6 times higher than sex-biased adult survival. Hatching sex ratio and mating system made negligible contributions (Fig. 2c).

### Consequences of ASR bias and polygamy on population viability

Biased ASR and polygamy create conditions whereby reduced survival of the limiting sex can compromise population viability, which has important implications for conservation. Our perturbation analysis showed that population growth was most sensitive to adult survival under all hypothetical scenarios of ASR and mating system (Fig. 3). Adult female survival elasticities were highest under scenarios of male-biased ASR. As expected, there was no sex-specific sensitivity of vital rates under an unbiased ASR and monogamous mating system. However, elasticity was highest for adult males under an unbiased ASR and polyandry.

**Figure 3.**
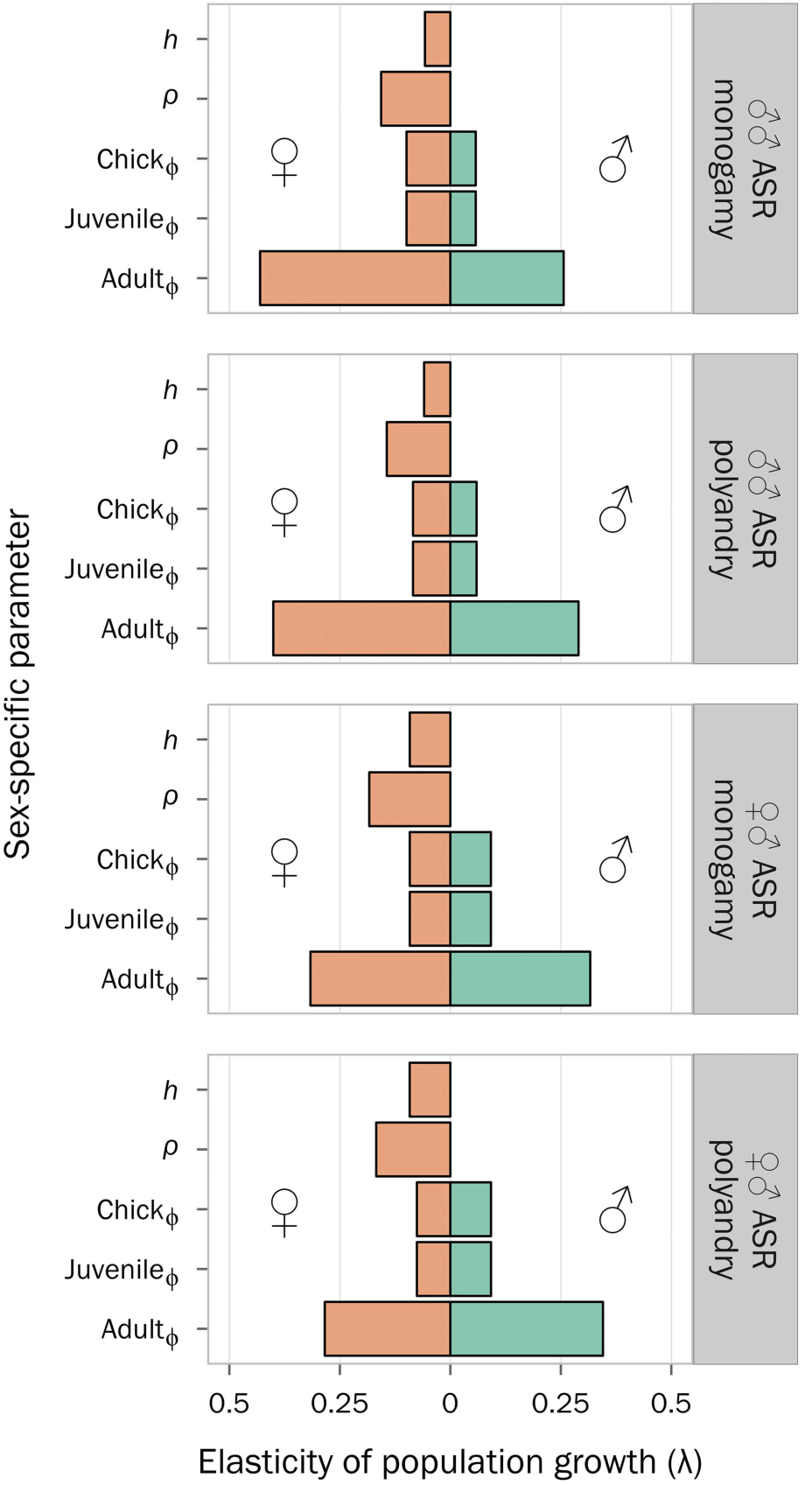
Sex-specific sensitivity analysis of population growth (λ) under four scenarios of ASR and mating system. Notation: *h* = mating system index, *ρ* = hatching sex ratio, ϕ = apparent survival.

To elucidate the conservation consequences of disregarding sex-biases, we compared the predictive accuracy of a detailed two-sex model incorporating polygamy to a conventional one-sex model. Over the seven-year study period, average population growth was below replacement (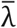_obs._ = 0.859 ± 0.28 SD, Fig. 4). This observed rate of decline was captured by the uncertainty distribution of the two-sex model (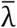_two-sex_= 0.849 [95% CI: 0.802−0.897], Fig. 4). In contrast, the one-sex version of the model exhibited greater uncertainty and significantly overestimated population growth (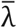_one-sex_ = 0.947 [95% CI: 0.883–1.01], Fig. 4).

**Figure 4.**
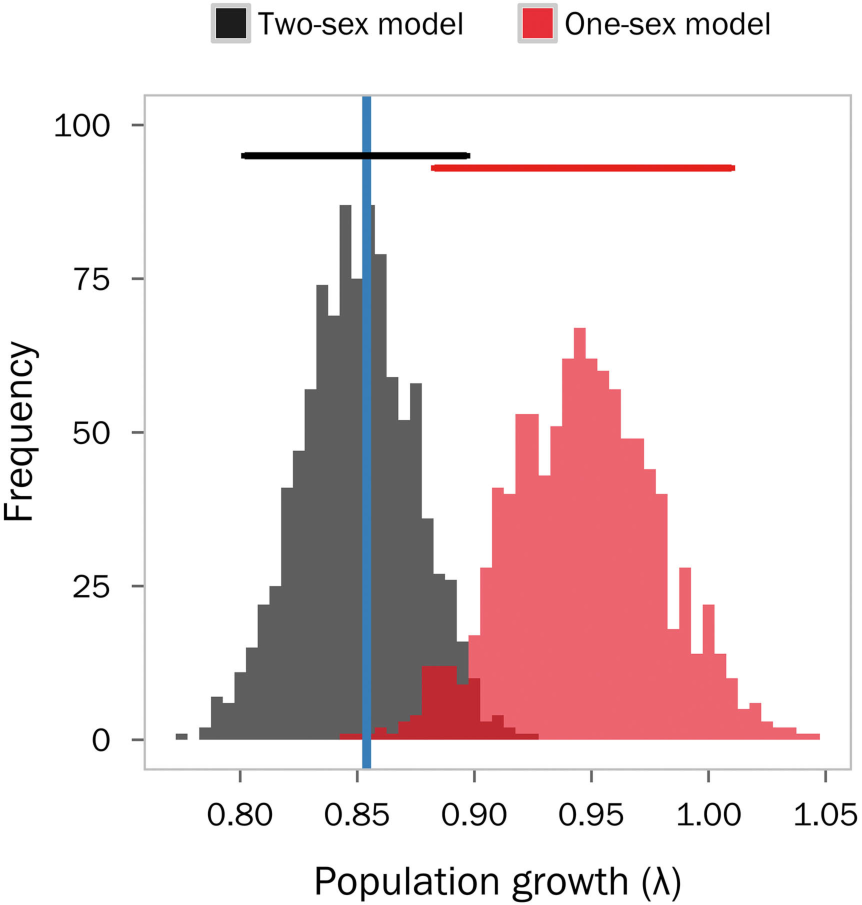
Distributions of deterministic population growth (λ) under a two-sex frequency-dependent matrix model (black) and a conventional one-sex matrix model (red) compared to the observed average annual population growth (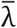) over the seven-year study period (blue vertical line). Horizontal bars above each distribution represent the 95% confidence intervals of the bootstrap simulation for each model.

## DISCUSSION

We present a comprehensive demographic model based on detailed individual-based life-history data from an intensively monitored bird population. By incorporating sex-specific feedbacks between survival and frequency-dependent reproduction, our model predicted a strongly male-biased ASR. This was complemented by our behavioral observations of a polyandrous mating system. Therefore, our findings build upon recent empirical and theoretical studies linking ASR to the evolutionary origins and consequences of mating system variation (10-12), and also provide novel insights into the sex- and stage-specific demographic components that contribute to ASR bias. Males had consistently higher apparent survival than females across all life stages, but a sex difference in the apparent survival of juveniles had the largest impact on ASR bias. Furthermore, population growth was most sensitive to perturbations in adult female survival under the male-biased ASR. Taken together, our results uncover the demographic pathways linking individual-level variation in survival and sex roles to population-level dynamics.

Obtaining reliable survival estimates from natural populations is challenging due to sex differences in behavior and life history. Our study addressed this uncertainty through mark-recapture models and bootstrapping. A central assumption of our model is that our marked subset of the population is representative of the entire population. This is appropriate given that we marked the vast majority of chicks and breeding adults in the population. Furthermore, we did not find a sex difference in the proportion of breeders versus nonbreeders (Fig. S1) indicating that our ASR estimate is not confounded by an excess number of unmarked non-breeding females.

Females had higher rates of polygamy than males, which is in line with most published records from other snowy plover populations (32, 35). This female-biased mating system complemented a strongly male-biased ASR. ASR in this species has previously been reported to be less extreme (34) than found here although the previous study was unable to incorporate sex-specific chick and juvenile survival, which made the greatest contributions to the ASR bias in this population (this study) and others (36). ASR bias is a widespread phenomenon in wild vertebrate populations, with mammals typically being female biased (mean ASR = 0.37 ± 0.15 SD) and birds typically being male biased (mean ASR = 0.55 ± 0.09 SD; Table S1; (37). Our ASR estimate for snowy plovers therefore lies within the natural variation observed in other avian taxa (9).

### Importance of early sex-specific demographic processes

Several hypotheses can be put forward to explain the observed pattern of male-biased early survival. Focusing first on the chicks, male hatchlings are significantly larger than their sisters in this population (38), potentially providing males with an advantage during early development. Another mutually non-exclusive possibility is that male chicks could achieve faster growth rates, as has been observed in Kentish plovers *Charadrius alexandrinus* (38), for example, either by virtue of sex-specific parental care (39) or as a consequence of sex-specific immunocompetence (40). Alternatively, predation could act sex-specifically, although male and female chicks do not differ appreciably in appearance and behavior, and we did not detect a sex difference in encounter rates (Fig. S2). Lastly, the sexes might differ in premature investment in sexual traits (41) although this seems unlikely as sexual ornamentation is moderate and body size differences are small (ref. 31; about 4%).

The sex bias during the juvenile stage made the largest contribution to the ASR bias. This corroborates the results of earlier avian studies showing that sex-specific first-year survival may lead to an ASR bias (25, 36). Our study goes further than previous works by decomposing the contributions from the first-year stage into chick and juvenile contributions. Juvenile survival contributed most towards ASR bias, probably because these naïve individuals face multiple challenges during the transition from parental independence to sexual maturity, potentially including predation, harsh winter climates and food shortages, all of which could disproportionately affect either sex. For example, in sexually size dimorphic species including red deer *Cervus elaphus* and great bustards *Otis tarda*, male young are less able to cope with severe winter weather (42) and food shortage (43), probably owing to the metabolic demands of large body size. In snowy plovers, such a mechanism is unlikely given the moderate size differences between males and females (31).

Another possibility is that sex-biased dispersal behavior could contribute towards sex-differences in apparent juvenile survival, as natal dispersal is typically female biased in birds (44), with snowy plovers being no exception (45, 46). However, dispersal and survival are not necessarily independent phenomena, as dispersal often entails survival costs such as increased predation risk and unfamiliarity of novel environments (47). Moreover, chicks are unable to disperse beyond the breeding site, so their survival estimate approximated true survival and thus implies a role of intrinsic sex differences in early survival. In addition, over the seven years of this study, few adults were resighted in adjacent populations and these individuals are unbiased with respect to sex. Finally, an independent study of snowy plovers in Monterey Bay, California found that survival was male biased even after accounting for sex-specific dispersal (34).

### Negligible effect of sex allocation

The hatching sex ratio, based on 340 hatchlings, was unbiased and served as a proxy for the primary sex ratio. Despite popular interest in sex-allocation theory (48, 49) relatively few studies have convincingly demonstrated offspring sex ratio biases in wild populations (50). Düsing (51), Fisher (52), and others (53-55) reasoned that if sex biases in survival emerged after the period of parental investment, sex allocation should not deviate from parity. This is precisely what we found, with ASR being strongly influenced by the sex-biased survival of independent juveniles, rather than deviations in the hatching sex ratio (Fig. 2). Furthermore, although the sex-biased survival of dependent chicks provided a noteworthy contribution to ASR bias (Fig. 2), fathers provide uniparental care of chicks in this species, and therefore the period of maternal investment typically ends at hatching. Given this parental care system, our result further confirms theoretical expectations of an unbiased hatching sex ratio.

### Evolutionary feedbacks between ASR and mating system

Mating systems are influenced by the availability of mates (11). A biased ASR creates conditions whereby one sex is in limited supply, thus forcing the other sex to compete for access to mates (7, 10). In shorebirds, ASR is a strong predictor of mating and parental strategies (13), with the limiting sex tending to have greater mating opportunities and reduced parental investment. These sex differences in the costs and benefits of parental care may facilitate polygamy (6).

However, the relationship between sex ratio and mating system represents a causality dilemma because of the positive feedback that polygamous mating systems impose on ASR bias and *vice versa* (56). On the one hand, polygamy entails sex-specific costs due to sexual selection which could drive the ASR bias, while on the other hand ASR bias creates uneven mating opportunities and thus facilitates polygamy. We show that sex biases originate prior to maturity (Fig. 5i), and are therefore likely influenced by natural selection. For example, genotype-sex interactions could impact chick survival during development (57) rather than during adulthood where sexual selection is expected to have a strong impact on survival. Consequently, ASR bias appears to drive the mating system rather than *vice versa* (Fig. 5ii). We cannot discount the possibility that sexual selection contributes towards sex biases in early survival, for instance via differential early investment in secondary sexual traits (41), but this seems unlikely as sexual dimorphism in plovers is negligible and, if anything, males are more ornamented than females (31, 58).

**Figure 5.**
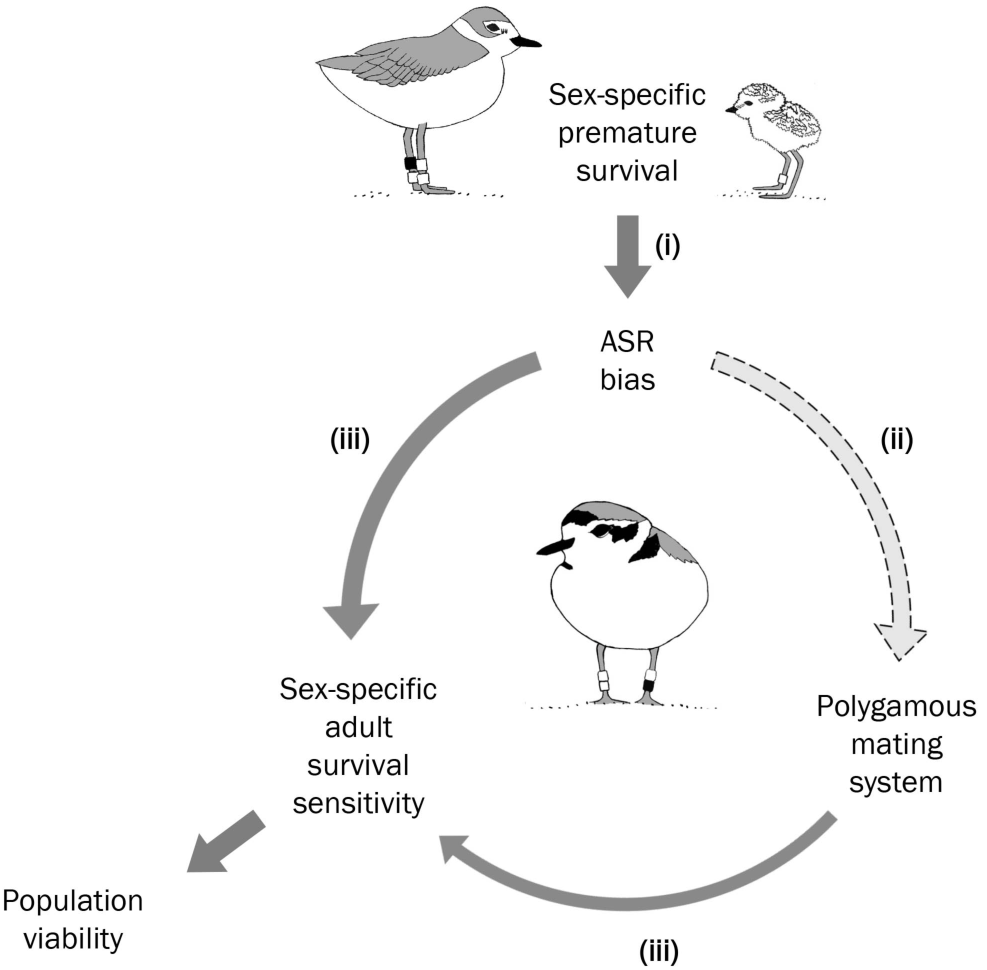
Schematic representation of the sex- and stage-specific demographic feedbacks between ASR, mating system, and population viability. In our study system, ASR bias is driven by sex-specific survival of early life-history stages (i), and likely facilitates a polygamous mating system (ii). Both, ASR bias and polygamy, contribute to sex-specific elasticities of population growth (iii), with ASR having the strongest effect as illustrated by arrow width.

### Consequences of mating system and ASR on population growth

In vertebrates, adult survival is often the most important parameter influencing population growth because adults have the greatest reproductive potential (59). This was the case in our population, but the effect of adult survival was sex-specific due to ASR bias and polygamy. Under a male-biased ASR, adult female survival had the highest elasticity to population growth, meaning that a small perturbation to adult female survival has a larger effect on population growth than an equivalent perturbation in all other parameters. Attempting to dissect the effects of ASR and mating system, we found that sex-specific elasticities of adults were greatest under scenarios of male-biased ASR compared to scenarios of polyandry (Fig. 5iii). Conversely, in scenarios of unbiased ASR, both sexes had the same elasticities for all parameters under monogamy, but under polyandry, adult male survival had the highest elasticity. These contrasts highlight the reproductive constraints imposed on populations with a biased ASR. When ASR is male-biased, polyandrous mating strategies optimize individual fecundity, and thus maximize population growth (1). However, reproduction is dependent on the availability of gametes, and thus a population with female-biased ASR and a polygynous mating system may not be limited in the same way because a single male can produce many more gametes than a single female and hence father more offspring. Therefore, similar studies of polygynous populations are needed to critically test the predictions of mating system, ASR, and population growth.

### Implications of two-sex vs one-sex modeling for biodiversity conservation

Modeling population trajectories is an important tool for conservation and management (60). Sex biases could potentially play an important role in population dynamics, particularly for polygamous species, yet sex-structured models remain uncommon. Here, we compare a conventional one-sex model to a two-sex model incorporating frequency-dependent reproduction and sex-specific survival. We show that our two-sex model provides a better fit to the observed data than a one-sex model. Moreover, while both models indicate negative population trajectories, the one-sex model substantially underestimates the rate of population decline and exhibits greater uncertainty. This has important conservation implications, especially for sex-biased populations of threatened or endangered species, where failing to incorporate sex-specific vital rates could lead to erroneous conclusions and overestimated population viability.

### Conclusions

With the rise of conservation-based monitoring of individually marked populations (61), a wealth of demographic data present new scientific horizons for ecologists and evolutionary biologists interested in sex-specific modeling. By combining extensive individual-based sex-and stage-specific vital rates and structured population models, we show that a male-biased ASR in a natural population is driven by male-biased survival of early life history stages (Fig. 5i). Our results indicate that ASR likely drives mating system (Fig. 5ii) although further experimental and/or comparative studies are needed to establish the causal link. Both ASR and mating system facilitate sex-specific sensitivities to population growth (Fig. 5iii), with ASR having the strongest effect. Male-biased survival in snowy plovers is consistent with recent comparative studies (62, 56) suggesting that many bird species may exhibit male-biased ASR (9, 37). Thus, our study makes an important contribution to understanding a widespread phenomenon in natural populations and highlights that ASR variation likely acts as an important catalyst of mating system dynamics and population viability.

## METHODS

### Field and laboratory methods

Over the seven-year study period, we collected mark-recapture and individual reproductive success data during daily surveys of the study site over the entire three-month breeding season that typically spanned from mid-April to mid-July. Plover chicks and adults were captured using funnel traps on broods or nests (63). We assigned adults a unique color combination of three darvic rings and an alpha-numeric metal ring, allowing the use of both captures and non-invasive resightings to estimate survival. Regular brood resightings combined with regular recaptures aided analyses of daily survival for chicks. Given our intensive nest search and capture efforts we are confident that we ringed the vast majority of chicks (>95%) and breeding adults (>85%) in the local population. Nests and broods were frequently monitored every two to seven days to assess daily survival and identify tending parents. During captures, approximately 25-50 μL of blood was sampled from the meta-tarsal vein of chicks and the brachial vein of adults for molecular sexing with the Z-002B marker (64) and verification with the Calex-31 marker located on the W chromosome (65). Details of our PCR conditions are found elsewhere (38).

### Quantifying mating system

We evaluated mating system of the population using a dataset that only included individuals for which we (i) were confident of the identity of their mates, and (ii) had observed them in at least two reproductive attempts that were either within the same season or in different seasons. Sex differences in the per capita number of annual mates were evaluated using a non-parametric Mann-Whitney-Wilcoxon test.

### Estimation of sex- and stage-specific survival

Our structured population model considered sex-specific survival across three key stage classes in avian life history: chicks, juveniles, and adults (Fig. 6). The chick stage was defined as the 25-day period between hatching and fledging during which offspring are dependent upon parental care (66). The juvenile stage was defined as the one-year transition period spanning from fledging to recruitment into the adult population. The adult stage represented a stasis stage in which individuals were annually retained in the population.

**Figure 6.**
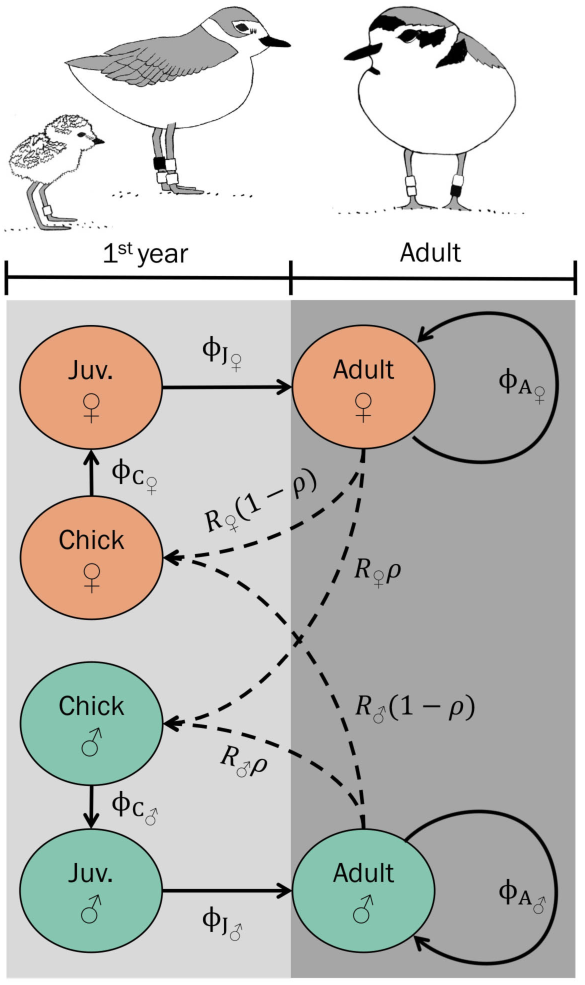
Snowy plover life cycle flow diagram illustrating sex-specific survival (ϕ) among three life stages (chick = *C*, juvenile = *J*, and adult = A) and the link between the sexes via the frequency-dependent mating function (*R*, see methods). The hatching sex ratio (*ρ*, proportion of male hatchlings) serves as a proxy for the primary sex ratio and allocates progeny to the male or female chick stage. Lower-level parameters (i.e. chick and juvenile survival) constitute the transition of first-year individuals illustrated in light grey.

We used mark-recapture models to account for sex, stage, and temporal variation in encounter (p) and apparent survival (ϕ) probabilities as they allow for imperfect detection of marked individuals during surveys and the inclusion of individuals with unknown fates (29). We use the term “apparent survival” as true mortality cannot be disentangled from permanent emigration in this framework (29). Furthermore, only a few nearby populations are regularly monitored and we have limited evidence that marked individuals disperse. See *SI Methods* for further details of our survival analysis.

### Matrix model structure

We built a two-sex pre-breeding Lefkovitch matrix model for the population that incorporated all three stages of plover life history into two annual transitions denoting first-years and adults (Fig. 6). Transitions of projection matrices are required to have equal temporal durations (67), and thus the chick stage (25 days) was combined with the juvenile stage (~11 months) as lower-level matrix elements to describe the transition of premature individuals into adulthood (Fig. 6). The projection of the matrix for one annual time step (*t*) is given by:

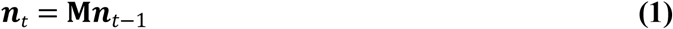

where ***n*** is a 4 × 1 vector of the population distributed across the two life stages and two sexes:

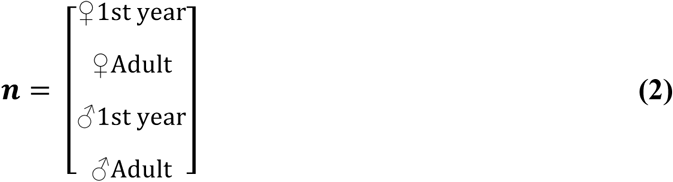

and **M** is expressed as a 4 × 4 matrix:

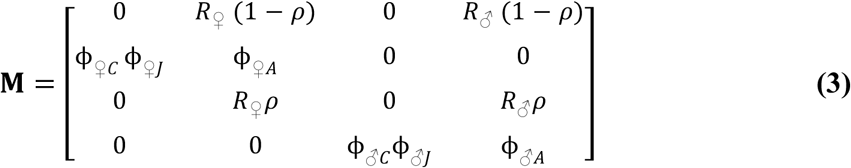

where transition probabilities (ϕ) between life stages are the survival of chicks (*C*), juveniles (*J*), and adults (A) for females (♀) and males (♂). The hatching sex ratio (*ρ*) describes the probability of hatchlings being either male (i.e. ρ) or female (i.e. 1 − ρ). Per-capita reproduction of females (R_♀_) and males (*R_♂_*) is expressed through sex-specific mating functions used to link the sexes and produce progeny for the following time step of the model given the relative frequencies of each sex (67). Here, we use the harmonic mean mating function which accounts for sex-specific frequency dependence (68):

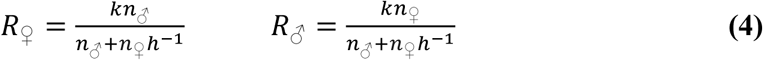

where *k* is the modal clutch size (3 in the case of snowy plovers), *h* is an index of the mating system (*h* > 1 signifies polygyny, *h* = 1 monogamy, and *h* < 1 polyandry), and *n_♀_* and *n_♂_* are the densities of females and males, respectively, in each time step of the model. In accordance with the predominantly polyandrous mating system, *h* was defined as the inverse of the average annual number of mates per female:

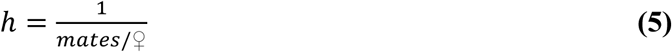

which we estimated from our behavioral observations of mating system (see above). To account for potential sex-biases arising prior to the chick stage (i.e. sex-allocation), we evaluated if the hatching sex ratio deviated significantly from parity (see *SI Methods* for details).

### Estimation of the adult sex ratio

We estimated ASR from the stable stage distribution (**w**) of the two-sex matrix model:

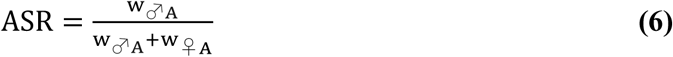

where w_♂A_ and w_♀A_ give the proportion of the population composed of adult males and females, respectively, at equilibrium. To evaluate uncertainty in our estimate of ASR due to sampling and process variation in our survival parameters, we implemented a bootstrapping procedure that resampled our mark-recapture data (see *SI Methods* for details).

### Life table response experiment of ASR contributions

Perturbation analyses provide important information about the relative effect that each component of a matrix model has on the population-level response, in our case ASR. To assess how influential a sex bias in parameters associated with each of the three life stages was on ASR dynamics, we employed a life-table response experiment (LTRE). A LTRE decomposes the difference in response between two or more “treatments” by weighting the difference in parameter values by the parameter’s contribution to the response (i.e. its sensitivity), and summing over all parameters (67). Here, we compared the observed scenario (**M**), to a hypothetical scenario (**M**_0_) whereby all female survival rates were set equal to the male rates and the hatching sex ratio was unbiased (i.e. *ρ* = 0.5). Thus, our LTRE identifies the drivers of ASR bias by decomposing the difference between the ASR predicted by our model and an unbiased ASR (25).

The contributions (C) of lower-level demographic parameters (θ) were calculated following Veran and Beissinger (25):

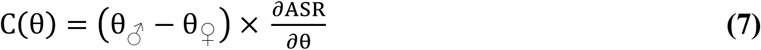

where 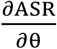 is the sensitivity of ASR to perturbations in the demographic rate θ in matrix **M**’, which is a reference matrix “midway” between the two scenarios (67):

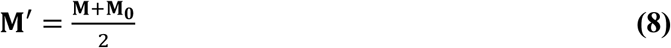

The two-sex mating function makes our model non-linear in the sense that the projection matrix, and specifically the fertility elements (Eq. 4), depend on sex-specific population structure. Perturbation analyses must therefore accommodate the indirect effects of parameter perturbations on population growth via their effects on population structure. To estimate the sensitivities of the vital rate parameters to ASR we employed numerical methods that independently perturbed each parameter of the matrix, simulated the model through 1000 time steps, and calculated ASR at equilibrium. This produced parameter-specific splines from which 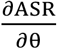 could be derived. Our approach appropriately accounts for the non-linear feedbacks between vital rates and population structure, though it does not isolate the contribution of this feedback (26, 69).

### Population growth consequences to ASR bias and a polygamous mating system

Biased ASR and polygamous mating systems can restrict the reproductive potential of a population due to a scarcity of the limiting sex (70). Thus, population viability can be indirectly affected by ASR and mating system via the sex-specific effects that vital rates have on population growth under a biased ASR or a polygamous mating system, or both (71). To investigate the relative influence that a biased ASR or a polygamous mating system has on population growth, we conducted a sensitivity analysis of all sex-specific parameters using four scenarios of the two-sex model: (i) polyandrous and male-biased ASR (i.e. the observed scenario), (ii) polyandrous and unbiased ASR, (iii) monogamous and male-biased ASR, and (iv) monogamous and unbiased ASR. In polyandrous scenarios, *h* was set to the value from field observations, whereas in monogamous scenarios, *h* = 1. In scenarios of unbiased ASR, male survival rates were assigned to both sexes (i.e. **M**_0_ above), whereas the original sex-specific structure was retained in male-biased scenarios.

Under each scenario, sensitivities of λ to perturbations in each parameter (θ) were estimated numerically as described above. Sensitivities were rescaled into elasticities (e), which describe the proportional response of λ to a proportional perturbation of a demographic parameter (67). This way, the sensitivity of parameters become directly comparable. Elasticities were calculated as:

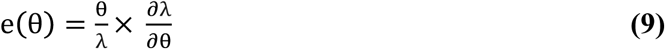

where 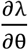 is the sensitivity of λ to perturbations in parameter θ.

### Comparison of two-sex versus one-sex models

Two-sex population models are rarely used in conservation biology because of the detailed data required to correctly parameterize them (70). As such, vital rates are typically estimated for only one sex or generalized across both sexes. However, in polygamous species, reproductive success varies according to the relative frequencies of mates (71), which is dictated by ASR and sex-specific survival. Therefore, ignoring sex-specific vital rates in polygamous species could misinform conservationists and wildlife management of population viability.

To explore how population growth varies under a two-sex model and a conventional one-sex model, we compared deterministic population growth of the two-sex model (**M**) to that of a one-sex model in which rates were averaged over both sexes (**A**):

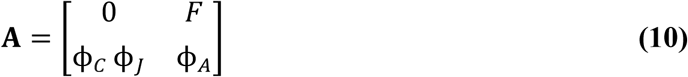

where *F* is the average annual per capita fecundity of females (expressed as hatchlings), and ϕ is the sex-averaged survival of chicks (C), juveniles (J), and adults (A). Deterministic growth (λ) was calculated as the dominant eigenvalue of **A** and as the asymptotic value of Σ w_t−1_/ Σ*w_t_* for **M**. To acknowledge uncertainty, we utilized the bootstrapped survival analysis described above by estimating the λ of each iteration under the structure of **A** or **M**. We contrasted the central tendency and spread of these distributions to one another and to the arithmetic average λ of the actual population trend over the seven-year study period.

All of our modelling and statistical analyses were conducted using R version “Bug in Your Hair” (72) with significance testing evaluated at α = 0.05. We provide all computer code and documentation as a PDF file written in Rmarkdown together with all the raw datasets needed to reproduce our modeling and analyses (*SI Dataset*).

## ACKNOWLEDGMENTS

Financial support of this study was provided by a Deutsche Forschungsgemeinschaft (DFG) doctoral fellowship awarded to LEP and supervised by OK, JH, and TS (GZ: KR 2089/9-1, AOBJ: 600454). We thank Sonoran Joint Venture, Tracy Aviary Conservation Fund, and CONACyT (Convocatoria de Investigación Científica Básica 2010, Grant no. 157570) for funding fieldwork and L. Lozano-Angulo and M. A. Serrano-Meneses for logistical support. KM was supported by a NERC GW4+ studentship NE/L002434/1. Molecular sexing was provided by the NERC molecular genetic facility at Sheffield (NBAF547, NBAF933, NBAF441). TS was supported by the Hungarian Science Foundation (NKFIH - K 116310) and was a Fellow at the Advanced Institute of Berlin. Original plover illustrations were kindly provided by A. Patrick. We are grateful for the helpful guidance and constructive thoughts kindly offered by S. Beissinger, D. Catlin, M. Galipaud, M. Jennions, P. Korsten, J. Laake, A. Potiek, A. Shah, and two anonymous reviewers.

